# Contact dependent suppression of *Clostridioides difficile* sporulation by enterococci requires the endocarditis and biofilm associated pilus

**DOI:** 10.64898/2026.04.22.718763

**Authors:** Alicia K. Wood, Caspar S. Carson, Holly R. Neubauer, Lesly-Hannah Gutierrez, Mofiyinfoluwa Adeoye, Angus Johnson, Ana Beatriz Garcez Buiatte, Bryce Chong, Laura C. Cook, Adam Session, Cheryl P. Andam, Peter T. McKenney

## Abstract

*Clostridioides difficile* is a healthcare-associated infection that arises when broad-spectrum antibiotic treatment disrupts the gut microbiota and is transmitted by highly resistant spores. Vancomycin-resistant *Enterococcus faecium* (VRE) is an opportunistic pathogen frequently co-isolated from *C. difficile* patients. We found that *C. difficile* sporulation is significantly reduced in VRE-*C. difficile* co-culture. Physical separation of *C. difficile* and VRE in transwell co-culture restored sporulation. Mixed macrocolony culture assays on solid agar confirmed physical contact is necessary for sporulation inhibition. We screened a panel of enterococci and found that most strains reduce sporulation, except *Enterococcus saccharolyticus*, which lacks predicted surface displayed virulence factors in its genome. We performed a candidate gene screen using an *Enterococcus faecalis* OG1RF transposon library and found that an insertion in the major pilin *ebpC* partially restored *C. difficile* sporulation in co-culture. These data were confirmed with in-frame deletions in the *ebpABC* pilus operon and a clinical isolate of *E. feacalis* lacking *ebpABC*. These findings suggest enterococci modulate *C. difficile* sporulation through a contact-dependent mechanism involving the Ebp pilus.

**Importance:** A characteristic of *C. difficile* infection is multiple episodes of acute disease. Spores are the transmission vector of *C. difficile* and are necessary for recurrence in models of disease. Our research demonstrates that *C. difficile* spore production is significantly reduced in the presence of enterococci, a common group of beneficial and pathogenic bacteria present in the gut microbiota. Physical contact with enterococci reduces *C. difficile* spore production. We attribute this effect to a protein structure on the surface of enterococci. This finding suggests a potential role for enterococci and the gut microbiota in general to uncover regulators of *C. difficile* spore formation. This may provide an avenue for innovative treatment strategies that reduce spore formation.

## Introduction

The gastrointestinal tract of mammals hosts a complex microbial ecosystem essential for nutrient processing, intestinal epithelial differentiation, and immune regulation (J.-Y. Lee, Tsolis, and Bäumler 2022). A diverse and healthy gut microbiota supports these functions as well as colonization resistance to bacterial infectious diseases (Woelfel, Silva, and Stecher 2024). Treatment with broad-spectrum antibiotics diminishes microbial diversity and increases susceptibility to *C. difficile* infections (CDI) (Buffie et al. 2012; Lawley et al. 2009). *C. difficile*, a principal etiological agent of nosocomial diarrhea, is an opportunistic, obligate anaerobe that produces metabolically dormant spores in its life cycle (C. D. Lee et al. 2022). Spores, which are intrinsically resistant to antimicrobials and most sanitizers used in hospitals, are crucial for CDI pathogenesis (Shen 2020) and transmission of infection (Deakin et al. 2012). Ingested *C. difficile* spores germinate in response to bile acids (Sorg and Sonenshein 2008) and grow into vegetative cells, causing infection in susceptible hosts. These active cells produce toxins that cause symptoms of CDI including diarrhea and potentially fatal pseudomembranous colitis (Kordus, Thomas, and Lacy 2022).

Vancomycin-resistant *Enterococcus faecium* (VRE) is commonly co-isolated from CDI patients (Tickler et al. 2020). Antibiotics treatment also results in expansion of VRE and its domination of the gut microbiota in humans and mice (Donskey et al. 2000; Ubeda et al. 2013). VRE domination increases the risk of both VRE bacteremia and subsequent *C. difficile* infection in stem cell transplant patients treated with antibiotics (Liao et al. 2021; Taur et al. 2012; Y. J. Lee et al. 2017). Enterococci are a core taxon of animal microbiota and have a high potential to modulate CDI virulence and the *C. difficile* life cycle. In co-infection mouse models, *C. difficile* virulence and toxin production were increased by both VRE and *E. faecalis* (Smith et al. 2022; Keith et al. 2020). Metabolic cross-feeding of arginine metabolites increased *C. difficile* virulence (Smith et al. 2022), while host-derived heme during active CDI increased levels of *E. faecalis* (Smith et al. 2024). Together these data suggest a reciprocal relationship between these two bacteria that have the potential to influence the severity of CDI.

Inter-species interactions also have the potential to affect sporulation. Entry into sporulation occurs during stationary phase and integrates nutrient sensing into the regulatory circuit that ultimately controls activation of the master regulator of sporulation Spo0A (Dineen, McBride, and Sonenshein 2010; C. D. Lee et al. 2022; Rizvi et al. 2023; Nawrocki et al. 2016; Antunes et al. 2012). In the densely populated colon, direct cell-to-cell contact with members of the gut microbiota also has the potential to modulate the *C. difficile* life cycle. Both *C. difficile* and enterococci genomes commonly encode pili (Kline et al. 2010; Ronish et al. 2024; Hancock, Murray, and Sillanpää 2014). Recently a type VII secretion system has been characterized in *E. faecalis* as well (Chatterjee et al. 2021, 2020).

Using a reductionist co-culture model, we found that co-culturing *C. difficile* and VRE resulted in a significant reduction in *C. difficile* sporulation, which is conserved among most enterococci. We confirmed that direct cell to cell contact was necessary for suppression of sporulation. Finally, using a candidate gene approach informed by comparative genomics, we identified the endocarditis and biofilm-associated pilus (Ebp) as necessary for mediating this suppression of sporulation.

## Results

### Enterococci inhibit *C. difficile* sporulation in co-culture

Previous work from our lab showed that in media containing an added fermentable carbon source, enterococci produce high amounts of organic acid that inhibit *C. difficile* growth (Neubauer et al. 2026). Here we optimized liquid co-culture in which both *C. difficile* VPI 10463 and *E. faecium* ATCC 700221 (VRE) achieved stable growth in protein-rich liquid sporulation medium (SMC) (Edwards, Suárez, and McBride 2013), which does not contain an added carbon source (Fig 1A-B). We found a significant reduction in heat-resistant spores and sporulation efficiency in *C. difficile*-VRE co-culture compared to *C. difficile* monoculture (Fig 1C-D) at 48 hours post inoculation. These data suggest that sporulation is inhibited in *C. difficile* – VRE co-culture under these conditions. To determine if sporulation is inhibited by VRE on agar, we performed macrocolony co-culture assays on agar plates. We found a significant decrease in sporulation under co-culture conditions, with no effect on total *C. difficile* or VRE CFUs (Fig1 D-E), however, heat-resistant spore production was significantly inhibited (Fig1 F-G).

**Figure 1:**
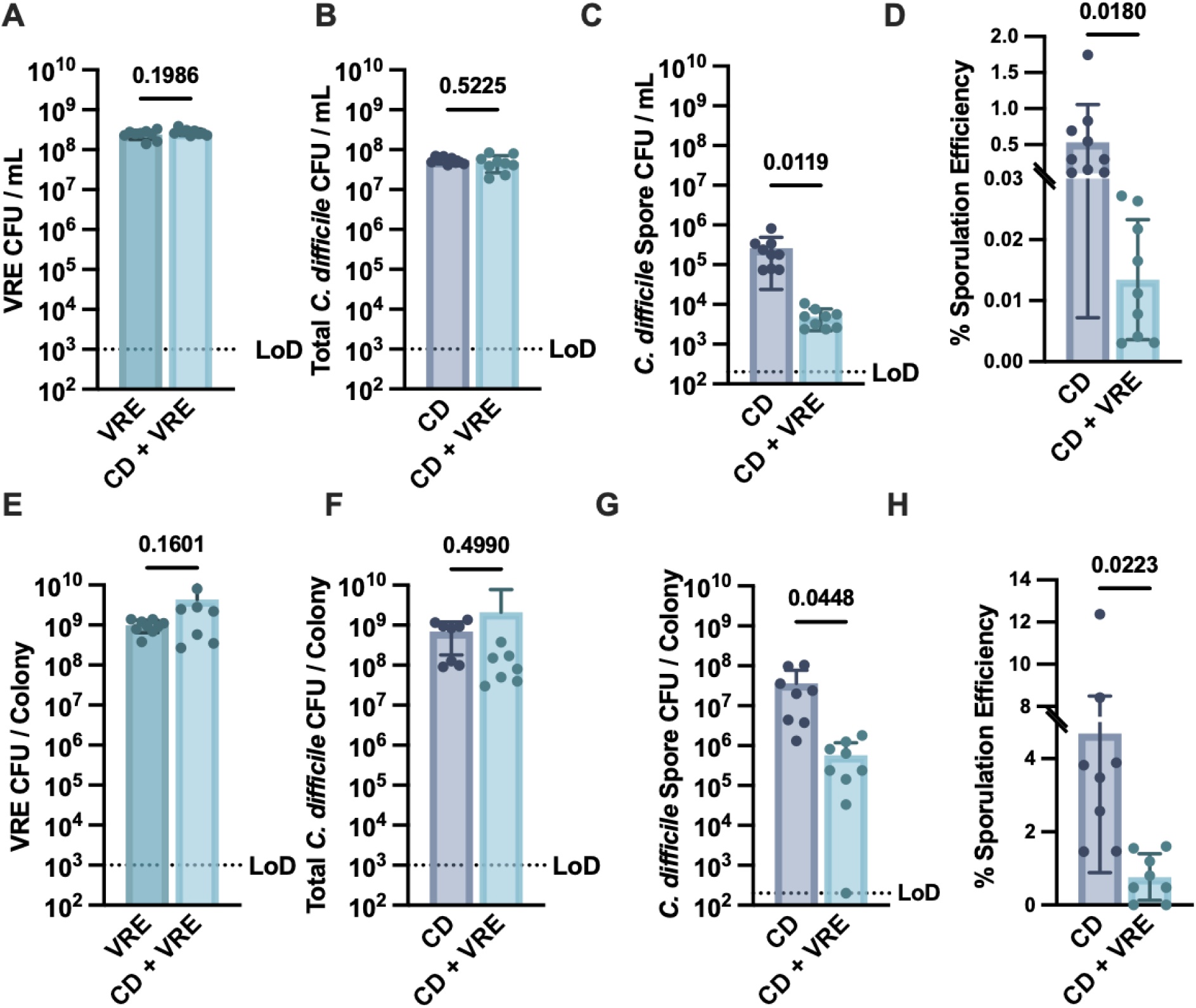
VRE Significantly Reduces *C. difficile* Sporulation in Coculture. (A-D) *C. difficile* and VRE cocultured anaerobically for 48 hours in liquid sporulation medium (SMC) in monoculture or co-culture followed by plating on selective media. (A) VRE CFUs. (B) Total *C. difficile* CFU (C) *C. difficile* heat-resistant spores. (D) Percent sporulation efficiency. (E-H) *C. difficile* and VRE cocultured anaerobically for 48 hours in mixed or single species macrocolonies on brain heart infusion (BHI) agar followed by plating on selective media. (E) VRE CFUs. (F) Total *C. difficile* CFUs. (G) *C. difficile* heat-resistant spores. (H) Percent sporulation efficiency. n = 8-9 biological replicates combined from 3 independent experiments. Statistics: Unparied t-test with Welch’s correction. LoD = Limit of Detection

To determine whether sporulation inhibition was conserved, we co-cultured *C. difficile* with a panel of enterococci that included commensal and pathogenic strains. Growth of enterococci and *C. difficile* were not significantly affected in co-culture (Fig2 A-B). Heat-treatment on coculture samples resulted in an ∼ 2-log reduction in spore CFUs and a significant reduction in sporulation efficiency in co-culture (Fig 2C-D). We found one strain, *Enterococcus saccharolyticus,* which is primarily isolated from non-clinical environments such as straw animal bedding (Rodrigues and Collins 1990) to be an exception (Fig 2E-H). *C. difficile* formed spores at levels comparable to monoculture when co-cultured with *E. saccharolyticus*.

**Figure 2.**
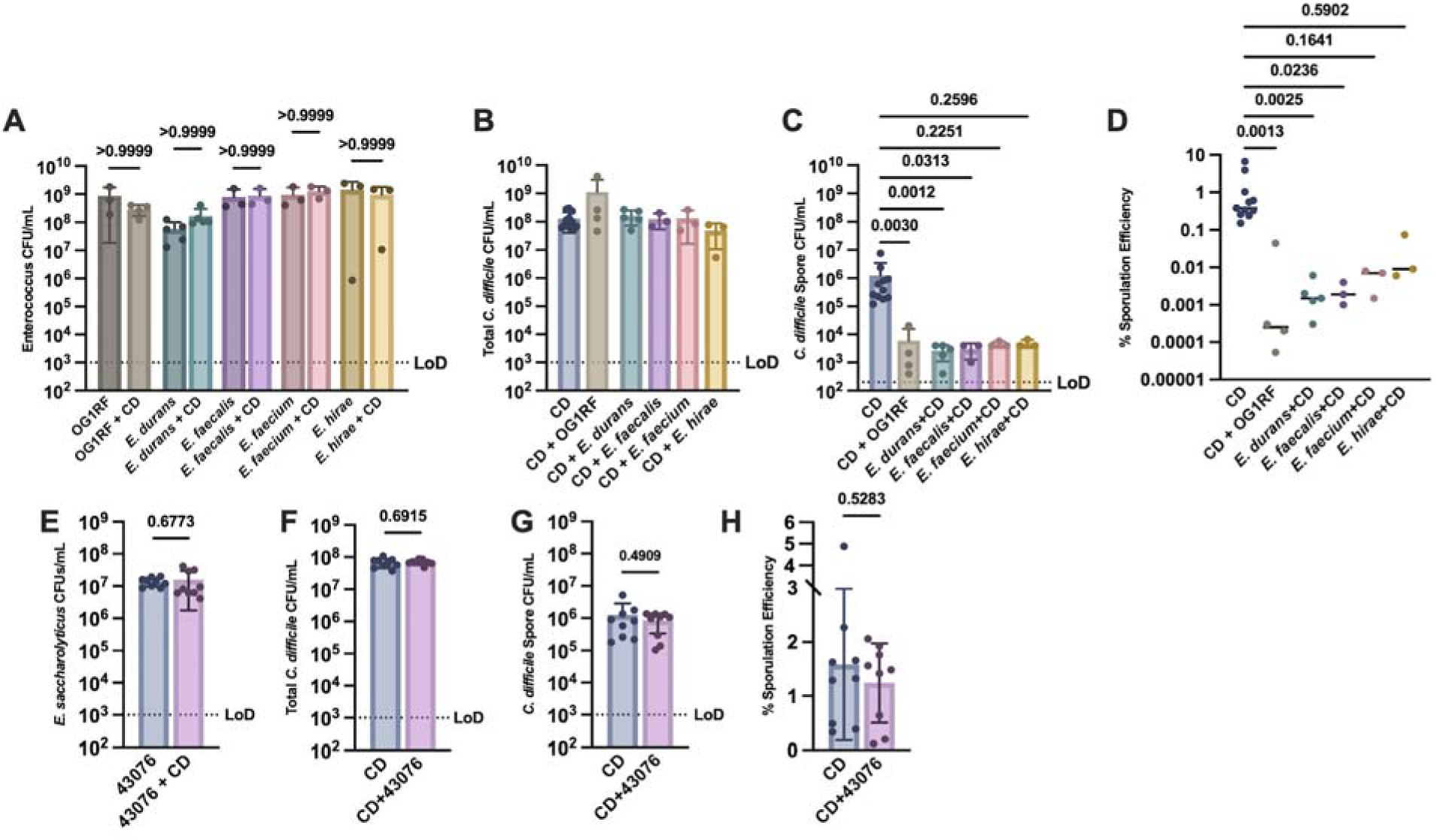
Inhibition of sporulation is conserved among enterococci: **(A-D)** Monoculture and coculture of *C. difficile* in liquid SMC anaerobically for 48 hours with *E. feacalis* OG1RF*, Enterococcus durans, Enterooccus faecalis, Enterococus faeicum,* or *Enterococus hirae* followed by plating on selective media. n = 3-11 biological replicates combined from 4 independent expeirments.**(A)** Enterococci CFUs **(B)** Total *C. difficile* CFUs **(C)** Heat-resistant *C. difficile* spore CFUs **(D)** *C. difficile* sporulation efficiency. **(E-H)** Monoculture and coculture of *Enterococcus saccharolyticus* and *C. difficile* in liquid SMC anerobically for 48 hours followed by plating on selective medium. n = 8-9 biological repicates combined from 3 independent experiments **(E)** *E. saccharolyticus* CFUs. **(F)** Total *C. difficile* CFUs. (**G)** Heat-resistant *C. difficile* spore CFUs. **(H)** Percent sporulation efficieincy. Statistics in A, E, F, G, H: t-test with Welch’s correction, statistics in B, C, D: Kruskal-Wallis One-way ANOVA with Dunn’s multiple comparisions test. LoD = Limit of Detection

We compared the genome of *E. saccarolyticus* with the other spore inhibitory enterococci and found approximately 200 genes predominantly associated with surface proteins and membrane processes that were absent. Notably, *E. saccharolyticus* lacks the conserved Endocarditis and biofilm-associated Pilus (Ebp) that is crucial for adhesion and biofilm formation in enterococci (Sillanpää et al. 2010; Nallapareddy et al. 2006, 2011; Kline et al. 2009). Our initial attempts to find homologs of the *ebp* pili in genomes of enterococci were limited to close relatives of *E. faecalis.* These predicted genes had low nucleotide and amino acid identity, as had been reported previously (Nallapareddy et al. 2011), suggesting a high level of heterogeneity. To search for pilus loci in a more sensitive manner, we searched a diverse collection of enterococci (Lebreton et al. 2017) for clusters of consecutive genes encoding predicted pilin domains (Fig 3). This analysis revealed two distinct families of pilus loci in the enterococci, orthologs of the canonical *ebp* cluster and a second family of more distantly related putative sortase-dependent pili. Many enterococci encode pili of both families. This search also failed to uncover a predicted pilus in the genome sequence of *E. saccharolyticus*, suggesting a potential role for pili in inhibition of sporulation.

**Figure 3.**
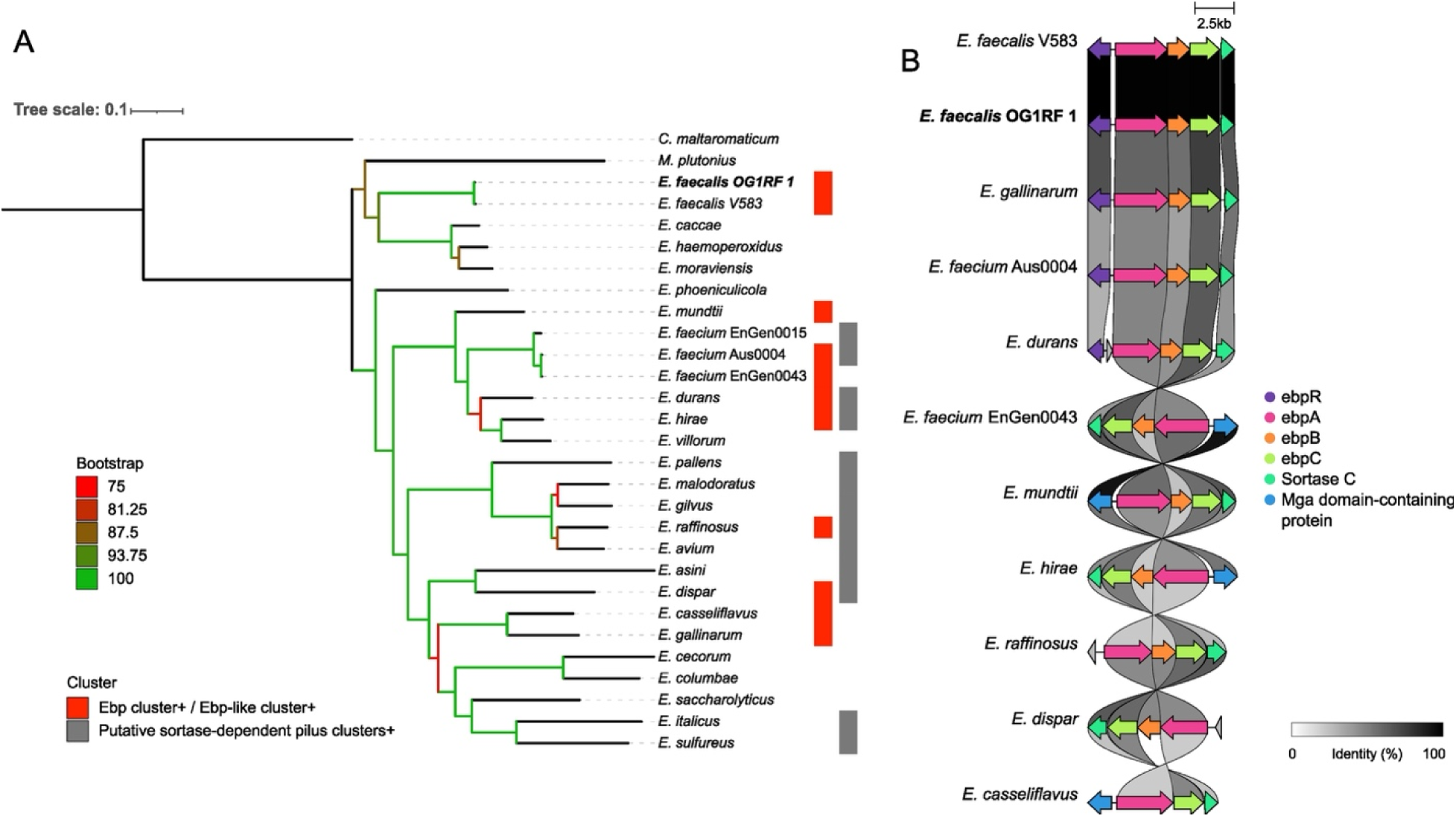
Multiple families of pili are present in enterococci. (A) Maximum likelihood phylogeny inferred from an alignment of 128 core genes. The tree scale indicates nucleotide substitution per site. *C. maltaromaticum* and *M. plutonius* were used as outgroups. Color strips indicate assemblies in which ebp/ebp-like clusters (red) or alternative putative sortase-dependent pilus clusters (grey) were identified. (B) Gene clusters aligned using Clinker. This figure shows genomes harboring ebp/ebp-like clusters that displayed detectable alignment to the reference locus (*E. faecalis* AE016830.1). Arrows represent annotated genes and are colored according to their functional class. Grayscale lines connecting homologous genes (arrows with the same colors) are shaded according to sequence identity.

To assess the conservation of the sporulation inhibition phenotype in multiple *C. difficile* strains, we co-cultured clinical isolate R20291 and lab derivative CD630Δ*erm* with VRE (Fig S1). Strain CD630Δ*erm* exhibited a not significant reduction in spore CFUs and a significant reduction in sporulation efficiency (Fig S1C-D). Strain R20291 exhibited a significant reduction in spore CFUs and no significant change in sporulation efficiency. We note that the effect size of spore CFUs in monoculture vs. co-culture appeared to be lower in these two strains (1-log or less) when compared to the data above from strain VPI10463 (generally ∼ 2-logs).

### Physical contact is required for inhibition of sporulation

Given the lack of pili and other surface structures in the genome of *E. saccharolyticus* (Fig 3), we hypothesized that cell to cell contact may be necessary for sporulation inhibition. To evaluate whether contact between strains was necessary, we optimized dual-species transwell co-culture, which allows free exchange of small metabolites through a permeable barrier but prevents direct cell contact. *C. difficile* exhibited wild-type sporulation levels when separated from VRE by transwell inserts (Fig 4). However, co-inoculation without inserts resulted in significant sporulation reduction, indicating physical contact is necessary for VRE to inhibit *C. difficile* sporulation (Fig 4C-D). We note that *C. difficile* growth and sporulation was higher in the transwell insert, when compared to the well of culture plate (Fig 4A-D). Presumably the transwell insert allows gas exchange with the surrounding atmosphere inside the culture plate.

**Figure 4.**
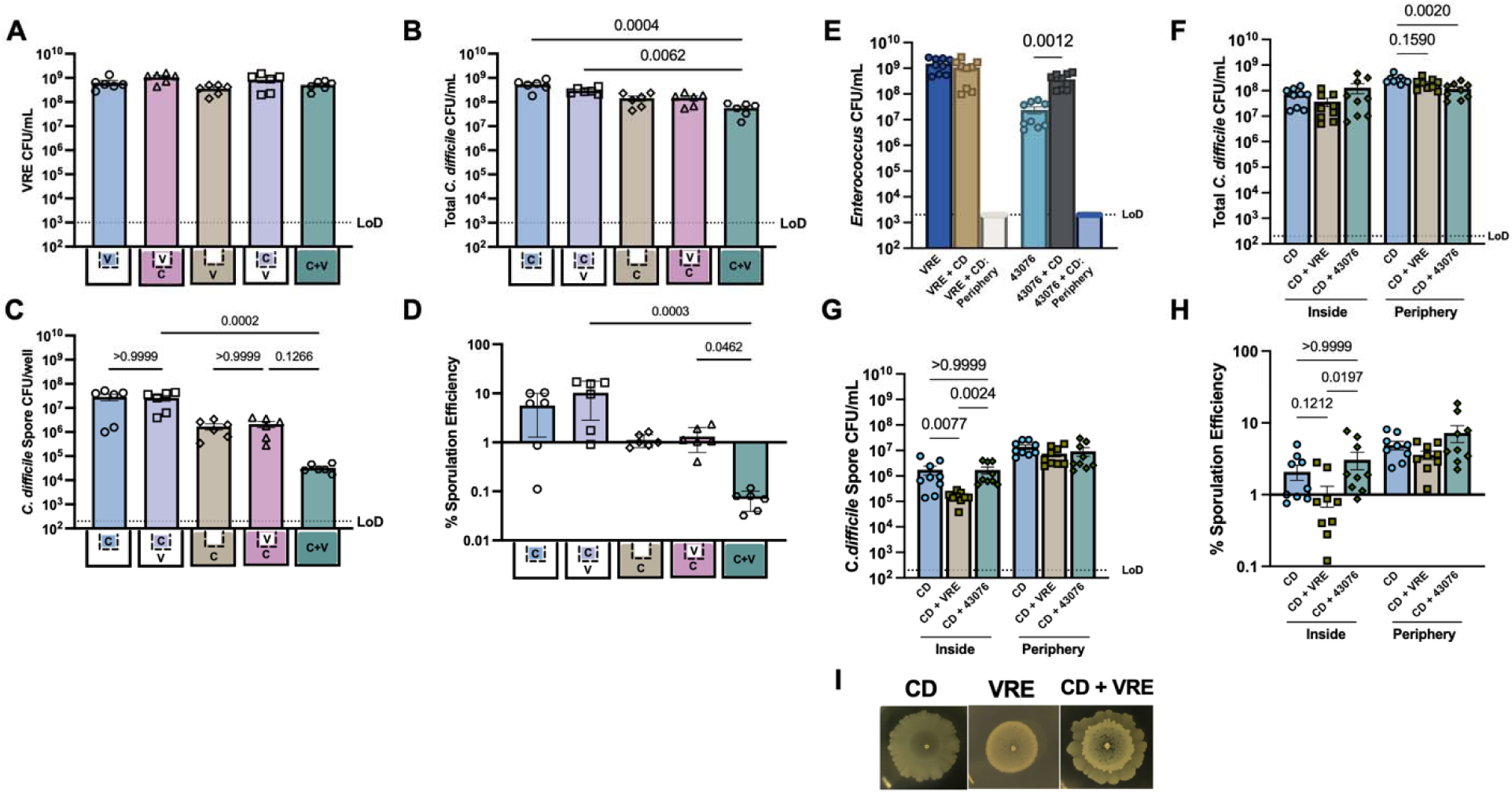
Sporulation inhibition is mediated by cell to cell contact. A-D) *C. difficile* and VRE were inoculated in transwells as indicated in the diagram on each x-axis. V = VRE, C = *C. difficile.* Transwells were incubated for 48 hours and plated on selective media. A) Total VRE CFUs. B) Total *C. difficile* CFUs. C) Heat resistant *C. difficile* spore CFUs. D) Percent sporulation efficiency. n = 6 biological replicates. Statistics: Pairwise comparisons for the effects of the presence and absence of a transwell: Unpaired Welch’s t-test. Multiple comparisons for differences in CFUs across conditions: Kruskall-Wallace One-way ANOVA with Dunn’s correction. E-G) Macrocolony migration assay, spots were inoculated on BHI agar and were incubated for 5 days followed by sampling the inside and periphery of the colony and selective plating. CD = *C. difficile*, 43076 = *E. saccarolyticus* E) Enterococcus CFUs, F) Total *C. difficile* CFUs, G) *C. difficile* heat resistant spore CFUs, H) Percent sporulation efficiency. Each data point represents one biological replicate combined from 3 experiments. Statistics: Kruskall-Wallace One-way ANOVA with Dunn’s correction. I) Representative images of 5-day single and dual species macrocolonies.

To further test the importance of cell-to-cell contact, we used a mixed macrocolony growth assay on solid agar. *C. difficile* is motile and forms rough colonies with the ability to spread from the inoculation point over time (Purcell et al. 2016). VRE is not motile and forms defined circular colonies when inoculated in a drop on a plate. This allowed us to set up a co-culture where *C. difficile* and VRE initially occupy the same space, from which *C. difficile* then migrates away. At the periphery, *C. difficile* escapes from and eventually loses contact with VRE (Fig 4E-I). In the middle of the microcolony *C. difficile* and non-motile VRE are mixed within the same space and maintain cell-to-cell contact. Sporulation on the periphery of a mixed macrocolony, reached levels similar to macrocolonies of *C. difficile* alone (Fig 4G-H). *C. difficile* sampled from the middle of the co-culture colony where direct contact with VRE is maintained had significantly reduced spore CFUs and sporulation efficiency (Fig 4G-H). When co-cultures of *E. saccarolyticus* and *C. difficile* were inoculated as macrocolonies, we did not detect significant changes in sporulation (Fig 4G-H). We do note a significant increase in *E. saccarolyticus* CFUs in the presence of *C. difficile* (Fig 4E), and a significant decrease in total *C. difficile* CFUs in the periphery of dual species macrocolonies (Fig 4F), suggesting an interaction between the two species in these conditions. Taken together, these data suggest that physical contact is required for enterococci to inhibit sporulation.

### Endocarditis and biofilm pilus (Ebp) is necessary for inhibition of sporulation

Given that cell surface components appeared to be missing from the *E. saccharolyticus* genome, we performed a limited candidate gene screen of an arrayed transposon library in *E. faecalis* OG1RF (Dale et al. 2018). We identified an insertion in the *ebpC* gene that significantly increased spore CFUs and sporulation efficiency when compared to co-culture with wild type OG1RF (Fig 5A-C). Location of the transposon insertion and validation of a single insertion was confirmed by PCR and whole genome sequencing (Figure S2). Previous deletion mutants in *ebpC* had reduced biofilm formation (Nallapareddy et al. 2006). The *ebpC::Tn* strain also had reduced biofilm formation (Figure S2), suggesting that the insertion is a loss of function. The gene *epbC* encodes the major pilin of the endocarditis and biofilm pilus (Ebp) (Nallapareddy et al. 2006; Nielsen et al. 2012). To further interrogate these findings, we assayed OG1RF deletion strains on agar, including single in-frame deletions of *ebpA*, *ebpB*, and *ebpC*, as well as a deletion of the entire *ebpABC* operon (Sillanpää et al. 2013). In this series of strains, only a complete deletion of the *ebpABC* operon did result in a significant increase in sporulation efficiency when compared to *C. difficile* co-cultured with wild type OG1RF (Fig 5D-F). We note that deletion of *ebpA*, the tip pilin, suppressed sporulation similar to OG1RF wild type, discussed in detail below. These data suggest that the Ebp pilus is necessary for suppression of *C. difficile* sporulation. The *ebp* locus is part of the core genome of *E. faecalis* (Nallapareddy, Sillanpää, et al. 2011). The clinical isolate *E. faecalis* strain TX1346, lacks the *ebpABC* operon (Nallapareddy et al. 2011) and also failed to suppress sporulation of *C. difficile* (Fig 5G-I), further demonstrating the importance of the Ebp pilus in this phenotype. Finally, we tested if pili were sufficient to inhibit sporulation by co-culturing *C. difficile* with wild type OG1RF that had been fixed with paraformaldehyde. Under these conditions the sporulation efficiency of *C. difficile* was not significantly changed when compared to monoculture (Fig 6), suggesting that pili are required on live cells of OG1RF to suppress sporulation. Taken together our data suggest that diverse enterococci inhibit *C. difficile* sporulation in a contact-dependent manner that requires the Ebp pilus on live cells.

**Figure 5.**
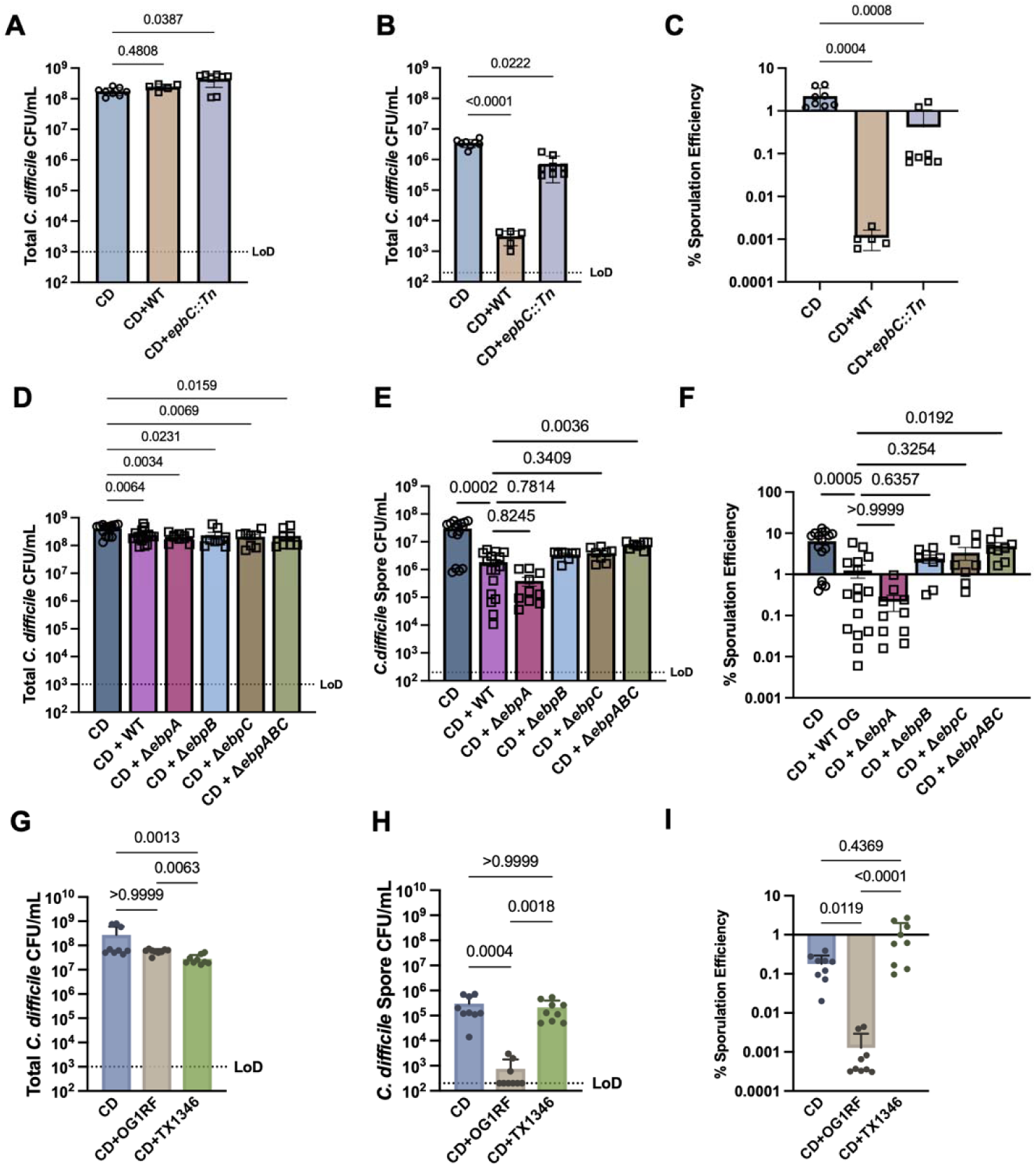
Sporulation inhibition requires the *ebp* pilus. A-C) The indicated strains were co-cultured in SMC liquid medium for 48 hours followed by heat treatment and selective plating. A) Total *C. difficile* CFUs, B) Heat resistant spore CFUs, C) Percent sporulation efficiency, n = 5-8 biological replicates per condition. D-F) The indicated strains were co-cultured on BHI agar for 48 hours followed by heat treatment and selective plating. D) Total *C. difficile* CFUs, E) *C. difficile* heat-resistant spores, F) Percent sporulation efficiency, n = 8-16 biological replicates per condition. G-I) The indicated strains were co-cultured in SMC liquid medium for 48 hours followed by heat treatment and selective plating. G) Total *C. difficile* CFUs, H) Heat resistant spore CFUs, I) Percent sporulation efficiency, n = 9 biological replicates per condition. All Statistics: Kruskal-Wallace One-way Anova with Dunn’s correction.

**Figure 6.**
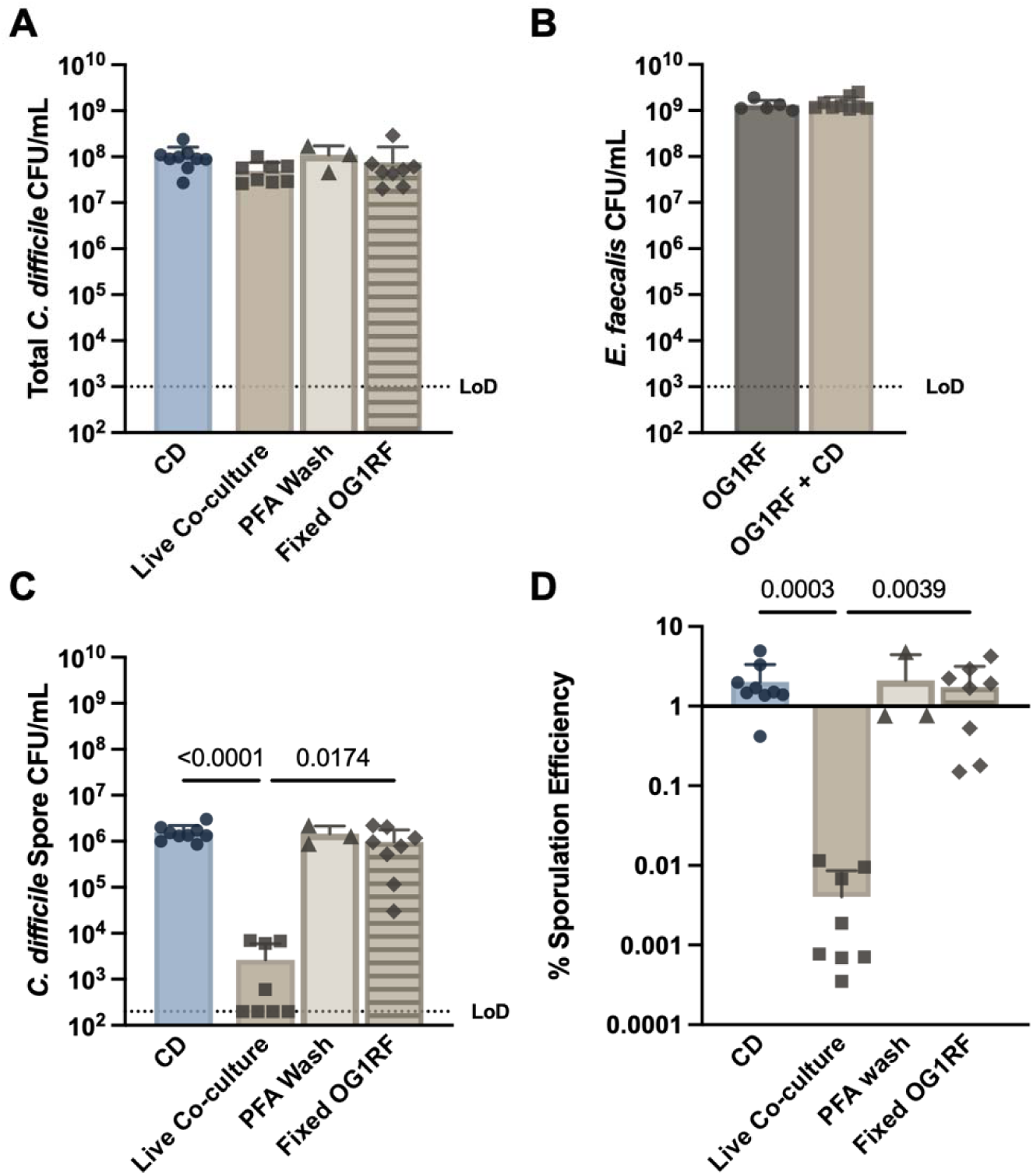
Live *E. faecalis* is required to suppress sporulation. *C. difficile* was cultured for 48 hours in liquid SMC with the indicated treatment: CD = equivalent volume sterile medium, Live Co-culture = live wild type OG1RF, PFA Wash = Mock treatment of sterile medium with PFA washes, Fixed OG1RF = wild type OG1RF fixed with 1% PFA. A) Total *C. difficile* CFUs, B) *E. faecalis* CFUs in Live Co-culture. C) *C. difficile* heat resistant spores. D) *C. difficile* sporulation efficiency. n = 3-9 biological replicates, statistics = Kruskal-Wallace One Way ANOVA with Dunn’s correction.

## Discussion

Here we describe an interspecies cell contact-mediated suppression of *C. difficile* endospore formation by enterococci that requires the endocarditis and biofilm-associated pilus. We identified a strain containing a transposon insertion in the major pilin gene *ebpC* that failed to suppress sporulation. Deletion of *epbABC* was sufficient to partially rescue sporulation efficiency, which suggests that other loci in OG1RF may be involved in mediating this phenomenon. Deletion of *ebpA,* which encodes the tip pilin, did not restore sporulation efficiency when compared to co-culture with wild type OG1RF (Fig 5). These results are intriguing as the Δ*ebpA* strain has been reported to produce abnormally long pili when imaged by immunogold transmission electron microscopy (Nielsen et al. 2012). Presumably, these pili are composed of long polymers of EbpC anchored to the cell wall by EbpB. It is possible that harvested OG1RF pili or engineered EbpC polymers may be sufficient to suppress sporulation, however our finding that fixed wild type OG1RF is unable to suppress sporulation (Fig 6) argues against this hypothesis. Together our data suggest that Ebp on live enterococci acts to suppress sporulation. To our knowledge this is the first report of a role for the Ebp pilus in interspecies bacteria-bacteria interactions. Pili have been proposed to mediate interspecies interactions in other systems. Interspecies electron transfer has been demonstrated between species of genus *Geobacter* and others by conductive e-pili (Lovley 2017).

Multiple groups have found that Ebp is expressed only on a sub-population of *E. faecalis* cells (Bourgogne, Thomson, and Murray 2010; Bourgogne et al. 2007; Nielsen et al. 2012; Kline et al. 2009; Afonina et al. 2018). Therefore, treatments and environmental conditions that alter the abundance of Ebp-positive *Enterococcus* cells in a population may in turn affect suppression of *C. difficile* sporulation. A potential physiologically relevant modulator of OG1RF piliation frequency is bicarbonate (Bourgogne, Thomson, and Murray 2010), which is secreted into the colon to regulate luminal pH (Becker and Seidler 2024). Full expression of the *ebpABC* locus requires the AtxA-family transcription factor EbpR, which is responsive to bicarbonate added to culture medium (Bourgogne, Thomson, and Murray 2010). Levels of the bicarbonate transporter *DRA* (*SLC26A3*) were decreased in the colon of *C. difficile* infected mice (Coffing et al. 2018; Peritore-Galve et al. 2023; Wang et al. 2020). Taken together we hypothesize that modulation of bicarbonate in response to CDI may affect *Enterococcus* piliation and in turn *C. difficile* sporulation during infection. Division of labor has been observed between sporulation and toxin production in vitro in populations of *C. difficile* (Donnelly et al. 2022). It is possible that inhibition of sporulation would result in increased toxin production and damage to the host.

In *E. faecalis* and *E. faecium*, deletion of the Ebp pilus has generally been associated with impaired biofilm formation in vitro (Nallapareddy et al. 2006; Sillanpää et al. 2010) and impaired colonization in some animal models of infection including models of endocarditis (Nallapareddy et al. 2006) and urogenital tract infections (UTIs) (Nallapareddy et al. 2011; Nielsen et al. 2012; Flores-Mireles et al. 2014). Deletion of *ebpABC* did not significantly alter colonization in the gastrointestinal tract in a chronic colonization model, despite a significant reduction in binding to mucus *in vitro* (Banla et al. 2019). There is some evidence that Ebp expression can lead to quantifiable differences in adherence properties of individual OG1RF cells. When OG1RF is exposed to magnetite and allowed to bind, followed by pull-down using a magnet, Ebp-positive cells are highly enriched in the magnetic fraction, suggesting that pili can mediate adherence heterogeneity within a population (Ho et al. 2025). Thus, it is possible that the Ebp-positive fraction of enterococci in our co-culture assays drive adherence to *C. difficile* sporangia and ultimately suppression of sporulation. Cellular heterogeneity also applies to the *C. difficile* side of this interaction, as co-culture of *E. faecalis* species resulted in an increase in *C. difficile* motility via CmrRST phase variation (Weiss et al. 2026). We note that the sporulation phenotype described here is conserved between *E. faecalis* and *E. faecium*, while modulation of phase variation was limited to *E. faecalis* (Weiss et al. 2026). This suggests some level independence between the regulation of the two phenotypes.

Contact-dependent inhibition systems were first characterized in Gram-negative bacteria. In *Escherichia coli,* the *cdiBAI* operon is sufficient for contact-dependent inhibition with CdiB localizing the toxin CdiA to the outer membrane while CdiI confers immunity (Hayes et al. 2014). Gram-positive bacteria contain a diverse family of LXG toxins that perform intra-species and inter-species contact-dependent inhibition via Type 7 secretion systems (Kobayashi 2021; Whitney et al. 2017). The OG1RF genome encodes a Type 7 secretion system and multiple potential LXG toxins (Chatterjee et al. 2021, 2020). It is possible that the Ebp pilus may facilitate close-range interaction between cells of OG1RF and *C. difficile* resulting in inhibition of sporulation or killing of sporangia via a secreted toxin. Both of these hypotheses are consistent with our data thus far. Distinguishing between inhibition of sporulation and the killing of sporangia would likely require anaerobic time lapse microscopy (Ribis et al. 2025) over a long time scale.

The contact-dependent suppression of sporulation characterized here is not mutually exclusive to metabolic influences between the two species (Smith et al. 2022). It is possible that short-range transfer of metabolites plays a role in the sporulation inhibition phenotype. Entry into sporulation in *C. difficile* is inhibited by glucose (Antunes et al. 2012), glycine (Rizvi et al. 2023), proline (Carter, O’Brien, and McBride 2025) and likely other nutrients. The activity of the master regulator of sporulation, Spo0A ultimately determines if a spore is formed (Deakin et al. 2012). A complex regulatory network integrates the nutritional state of the environment with Spo0A-P levels to determine the fate of an individual *C. difficile* cell (C. D. Lee et al. 2022). For example, at the transcriptional level the transcription factor CodY senses branched chain amino acids and GTP (Dineen et al. 2007) and represses genes involved in promoting sporulation (Dineen, McBride, and Sonenshein 2010). As levels of GTP in the cell fall, CodY dissociates from promoters and entry into sporulation proceeds (Dineen, McBride, and Sonenshein 2010; Nawrocki et al. 2016). CodY target genes also contribute to heterogeneity in sporulation efficiency between strains of *C. difficile* (Monteiro et al. 2026). Metabolism that depletes GTP levels in *C. difficile*, such as the synthesis of c-di-GMP (Dhungel and Govind 2021; Bordeleau et al. 2011; Purcell et al. 2012; Edwards et al. 2021) has the potential to affect sporulation. Over-expression of the di-guanylate cyclase DccA reduced sporulation in *C. difficile* in a manner that is dependent on an as yet unidentified modulator of sporulation efficiency (Edwards et al. 2021). Production of the alarmone (pp)pGpp (Pokhrel et al. 2020; Poudel et al. 2022) by *C. difficile*, or the consumption of GTP precursors or branched chain amino acids by enterococci also has the potential to affect initiation of sporulation.

Inter-species signaling and cell-to-cell contact among bacteria may be an underappreciated aspect of developmental cell fate decisions in spore forming Firmicutes (Bacillota). In the endospore former *Bacillus subtilis*, several inter-species interactions have been characterized that modulate sporulation. Conditioned medium of *E. coli* promoted sporulation (Grandchamp, Caro, and Shank 2017). A screen of the Keio knockout collection in *E. coli* identified 115 loci that were necessary for sporulation in co-culture with *B. subtilis*, including the siderophore enterobactin which was sufficient to promote *B. subtilis* sporulation (Grandchamp, Caro, and Shank 2017). Conversely, the siderophore coelichelin produced by environmental *Streptomyces* isolates inhibited sporulation which led to sensitization to the phage SPO1 (Zang et al. 2025). Additionally, a co-culture screen identified that sub-inhibitory concentrations of the broad spectrum antibiotic 2,4-diacetylphloroglucinol synthesized by pseudomonads reduced sporulation (Powers et al. 2015).

The sporangium producing organism itself may also compete with its neighboring species. *B. subtilis* sporulation itself was necessary to prevent predation by *Myxococcus xanthus* (Müller et al. 2014, 2015). In this model, secretion of the secondary metabolite bacillaene inhibited *M. xanthus*, allowing sufficient time for *B. subtilis* sporulation to complete. In mixed species co-culture with *Streptomyces coelicolor*, *B. subtilis* produced surfactin which promoted its own sporulation while limiting aerial hyphae and spore formation in *S. coelicolor* (Straight, Willey, and Kolter 2006). These data suggest that competitive interactions that affect sporulation may occur on both sides of the interaction between a spore former and another species.

We have described here that *C. difficile* endospore formation can be actively modulated by other microbes within a shared niche. It is possible that domination of the gut microbiota by enterococci, as occurs in humans during stem cell transplantation (Taur et al. 2012), may limit sporulation and increase production of toxin by *C. difficile.* An increase in *C. difficile* toxin production has been observed in mouse models of co-infection (Keith et al. 2020; Smith et al. 2022). We have not tested commensal clostridia for the effects of enterococci on sporulation. Commensal clostridia are generally associated with health and are components of FDA approved microbiota derived treatments for *C. difficile* (Feuerstadt et al. 2022; C. Lee et al. 2023). If sporulation is also required for engraftment and stable colonization of the intestine by commensal clostridia, it is possible that enterococci may modulate the effectiveness of such microbiota derived therapeutics as well.

## Acknowledgments

We thank Scott Hultgren, Cris Gualberto, Kavindra Singh, Karen Jacques-Palaz, Cesar Arias, Barbara Murray, Aimee Shen, Revathi Govind, Gary Dunny, Julia Willet, Breck Duerkop, Kimberly Kline, Eric Pamer, Alexander Rudensky, John Hambor and Su-Ellen Brown for strains and reagents. We thank members of the Binghamton Biofilm Research Center for comments on the project and support. This work was supported by NIH R21AI171634 to PTM and R03AI180614 to PTM and AS, R35GM142924 to CPA as well as startup funds to PTM from Binghamton University. AKW was supported by the Clifford D. Clark Fellowship from Binghamton University. MA and L-HG were supported by the McNair Scholars program at Binghamton University.

## Author Contributions

AKW performed all wet lab experiments with assistance from HRN, CRC, MA and L-HG. PTM and AJ performed pilot experiments that uncovered the initial phenotype. AS performed resequencing read mapping of the *ebpC::Tn* mutant. LCC assisted with OG1RF experimental design. ABGB, BC and CPA performed pilus gene cluster searches in enterococci. PTM and AKW wrote the manuscript with input from all co-authors.

### Disclosure

PTM is a co-inventor on US Patents 10,646,520, 11,207,374 and 11,471,495 owned by Memorial Sloan Kettering Cancer Center and receives licensing royalties originating from Seres Therapeutics Inc. and Nestle Health Sciences.

## Materials and Methods

### Bacterial strains and culture media

Strains are listed in Table 1. *C. difficile* strains were cultured in Brain-heart-infusion broth (HiMedia) supplemented with 0.5% yeast, 0.1% taurocholate, and 0.1% L-cysteine (BHIS-TA). Enterococci strains were cultured in BHI + 0.1% L-cysteine (BHIS). The wild-type OG1RF strain was cultured overnight in BHIS + 25ug/mL fusidic acid and 50ug/mL rifampicin. OG1RF transposon strains were cultured overnight in BHIS+10ug/mL chloramphenicol. OG1RF strains were grown in a shaking incubator aerobically at 37°C, shaking at 200rpm. For selective plating, *C. difficile* strains were enumerated on either BHI agar supplemented with 0.1% L-cysteine, 0.1% taurocholate, 16ug/mL cefoxitin, and 250ug/mL D-cycloserine (BHIS-CC-TA) or Cefoxitin-Cycloserine-Fructose Agar plates supplemented with 16ug/mL cefoxitin, 250ug/mL D-cycloserine, 0.1% cysteine, 0.1% taurocholate. All enterococci except OG1RF transposons were selectively plated on Pfizer Selective *Enterococcus* Agar (HiMedia). OG1RF transposons were plated on BHIS agar + 10ug/mL chloramphenicol. For biofilm assays OG1RF and transposons were cultivated in tryptic soy broth + 0.25% glucose. All experiments where *C. difficile* was cultivated were performed in an anaerobic chamber (Coy Laboratory Products) with an atmosphere of 90% N_2_, 5% CO_2_, and 5% H_2_.

### VRE-*C. difficile* coculture assay

Both *C. difficile* and VRE were grown to exponential phase (OD600 0.5-0.8) and normalized to an OD of 0.50. *C. difficile* and VRE monocultures were grown in parallel as controls to assess viable co-culture growth and as a control for *C. difficile* sporulation. Monocultures and cocultures were cultivated in 4 mL of sporulation media (SMC). All cultures were inoculated at ratio of 1 CD to 1 VRE or other *Enterococcus*. Samples were incubated for 48 hours at 37°C in the anaerobic chamber unless otherwise stated. For total CFU/mL enumeration, samples were serial diluted 1:10 in PBS and drop-pated on selective media. Plates were incubated for 24 hours before enumeration. For co-culture assays involving *E. feacalis* OG1RF an inoculation ratio of 200 CD to 1 OG1RF was used.

### Heat resistance assay

All samples containing *C. difficile* were heat-treated for spore CFU/mL enumeration in a water bath set to 65°C, confirmed by an additional thermometer. Samples were loaded into 0.2mL BioRad PCR strips and diluted 1:1 with PBS. The total treatment time was 30 minutes, but samples were gently mixed by pipetting up and down at 15 minutes to mitigate sample settling at the bottom of the tube. Undiluted and serially diluted samples were drop-plated on either BHIS-CC-TA or CCFA agar plates. Plates were incubated for 24-36 hours before enumeration. Sporulation efficiency was calculated as spore cfu/mLTotal CFU/mL ×100.

### Dual-species transwell assay

Dual-species transwell assays were set up in 24-well plates using 0.4um PET inserts (CellTreat). 1:10, 15mL SMC inoculums were made from OD-normalized exponential phase cultures to inoculate the wells and inserts. Autoclave-sterilized forceps were used to manipulate the transwell inserts. The well beneath the insert was filled with either 1mL of inoculum or 1mL of sterile SMC. Inserts were filled with either 250uL of inoculum or 250uL of sterile SMC. The surrounding wells were filled with PBS to prevent evaporation of the sample inoculated in the insert. The controls for the transwell assay were as follows: a barrier control where the insert was inoculated, and the well was filled with sterile SMC; a well control where the insert contained sterile SMC and the well was inoculated; and a “contact” control where no insert was present. Samples inoculated in the well were plated on both BHI-CC-TA and Pfizer Selective *Enterococcus* Agar to confirm no bacteria migrated through the insert filter. Disposable inoculation loops (Fisher) were used to harvest samples from the inserts and wells. The loops were used to remove any sample adhered to the filter or bottom of the 24-well plate. Subsequently, a pipette was used to gently mix and aliquot the sample into a sterile microfuge tube. The samples were vortexed on high for 2 minutes before being serially diluted and plated for total CFU/mL and heat-treated for spore CFU/mL enumeration.

### Co-culture macrocolony assays

To maintain consistency with no risk of sample overlap, coculture spot assays were performed in 12-well plates. Briefly, using a serological pipet, 3mL BHIS agar was pipetted into each well of a 12-well plate. Exponential-phase cultures were OD-normalized to 0.50 were loaded into a 96-well plate. The inoculums for this assay were optimized for stable co-culture growth on agar medium with an inoculation ratio of 1 VRE to 10 CD. The VRE monoculture control was 1 VRE to 10 sterile BHIS; CD monoculture control was 1 sterile BHIS to 10 CD. A spot of 10uL of sample was placed in the middle of each well. Plates were parafilmed, inverted, and incubated at 37°C for 48 hours before harvesting. The well plates were tilted 45° and flooded with 600uL of PBS to harvest the samples. Disposable inoculation loops were used to gently scrape the sample from the surface of the agar into the PBS. The sample was pipetted off the plate and aliquoted into a sterile microfuge tube. Samples were mixed by pipette before being vortexed on high for 2 minutes to help break up aggregates. Fully resuspended samples were serially diluted and plated for total CFU/mL and heat-treated for spore CFU/mL enumeration. For Inside / Periphery plating (Fig 4 E-I) 6-well plates were inoculated with 20uL of the appropriate monoculture or dual culture and were incubated for 5 days before harvesting. A sterile loop was used to harvest the periphery and inside of the macrocolonies followed by selective plating.

### Biofilm attachment assays and crystal violet staining

Biofilms were cultivated in tissue-culture treated, polystyrene 24-well plates (Greiner Bio-One). Exponential-phase cultures were OD-normalized to 0.50 with fresh media and inoculated 1:10 in tryptic soy broth + 0.25% glucose. Biofilms were grown aerobically and static at 37°C and processed either 2 hours to assess primary attachment or 24 hours to assess irreversible attachment. The bulk liquid was pipetted off and the biofilms washed 2X with PBS. Wells were fixed with 250uL of 4% PFA for 10 minutes and rinsed 2X with sterile DI water. Wells were dry before adding 250uL of 0.1% crystal violet for 15 minutes. The stain was pipetted off and wells were washed with sterile DI water until washes were clear. Wells were eluted with 80Ethanol:20Acetone solution and read at OD595 using a Beckman-Coulter DTX880 plate reader.

### PCR genotyping and whole genome sequencing of *ebpC::Tn*

Genomic DNA was extracted from wild-type OG1RF and *ebpC*::*Tn* using a Wizard Promega kit following manufacturer instructions. The concentration and purity of the DNA were assessed using a NanoDrop One spectrophotometer. PCR was performed to amplify the target gene *ebpC* using the following primers that flank the *ebpABCsrtC* locus. The PCR conditions were optimized to accommodate the 2.1kb transposon insertion. Wild-type *E. faecalis* strain OG1RF served as a positive control to verify the efficiency of the PCR and the transposon insertion. PCR products were visualized by agarose gel electrophoresis.

Illumina sequencing libraries of genomic DNA from *ebpC::Tn* was were prepared using the tagmentation-based and PCR-based Illumina DNA Prep kit and custom IDT 10bp unique dual indices (UDI) with a target insert size of 280 bp. No additional DNA fragmentation or size selection steps were performed. Illumina sequencing was performed on an Illumina NovaSeq X Plus sequencer in one or more multiplexed shared-flow-cell runs, producing 2×151bp paired-end reads. Demultiplexing, quality control and adapter trimming was performed with bcl-convert1 (v4.2.4). Reads were deposited at NCBI Short Read Archive under BioProject PRJNA1454558. The NC_017316.1 genome fasta and ASM17257v2 .gff files were downloaded from NCBI refseq. We downloaded the transposon sequence (Dale et al. 2018) and aligned to each assembly using BLASTN with an e value cutoff of 1e-2 (Camacho et al. 2009). Whole-genome alignment between the *ebpC::Tn* mutant and NC_017316.1 was done using minimap2 with default parameters (Li 2018). No alignments were found in the NC_017316.1 reference. A single alignment was found in the mutant genome. Alignments and annotation were visualized using IGV (Robinson et al. 2011).

### Genome annotation and phylogenetic reconstruction

The complete list of genomes, outgroups, references, and corresponding GenBank accession numbers used in this study is provided in Supplementary Table S2. Genome assemblies were retrieved from National Center for Biotechnology Information (NCBI) in FASTA format and reannotated *de novo* using Bakta v.1.11 (Schwengers et al. 2021) to ensure consistent and up-to-date functional annotations across all assemblies, and to generate uniform locus tag identifiers required for downstream analyses. All genome assemblies used in this study were evaluated for genome size, N50, and GC content using QUAST v.5.0.2 (Gurevich et al. 2013). The totality of genes in the dataset was determined with Panaroo v.5.2 (Tonkin-Hill et al. 2020), using the -strict flag to ensure that only high-quality sequences were included. Core genes were defined as those present in ≥95% of the genomes, whereas accessory genes were those detected in <95% of genomes. Nucleotide sequences for individual gene families were aligned using MAFFT (Katoh et al. 2002). A concatenated alignment of 128 core genes was generated and used to infer a maximum-likelihood phylogeny with IQ-TREE v.2.4 (Nguyen et al. 2015). Model selection was performed using ModelFinder (Kalyaanamoorthy et al. 2017), and the best-fit substitution model was GTR+F+I+R5. Branch support was assessed with 1,000 ultrafast bootstrap replicates (Hoang et al. 2018). We used iTOL (Letunic and Bork 2024) to visualize and annotate the phylogenetic tree.

### *In silico* identification of Ebp pilus and Sortase-dependent pilus clusters

To identify Ebp and putative sortase-dependent pilus clusters, we conducted tBLASTn (Gertz et al. 2006) searches using the protein sequences of EbpA, EbpB, EbpC and SrtC from the *E. faecalis* reference genome (NCBI accession no. AE016830.1) as queries against our genome assemblies. Given the phylogenetic diversity of the *Enterococcus* species included, a relaxed threshold of ≥20% amino acid identity and E-value < 1×10⁻^5^ were applied to maximize the detection of divergent homologs. In parallel, we scanned all genomes with hmmscan (HMMER v3.4) (Finn, Clements, and Eddy 2011) using the Pfam-A.hmm database (Sonnhammer et al. 1998) in default mode. Manual inspection was then performed to identify genomic regions containing at least one sortase flanked by a minimum of two proteins harboring pilin-associated domains, as defined by Pfam domain annotations. Hits with scores ≥30 and i-Evalue < 1×10⁻^5^ were retained for further analysis. When multiple domain matches for a given locus tag met the score and i-Evalue thresholds, only the highest-scoring hit was retained to assign a putative function to each locus tag for downstream genomic region extraction. Regions of interest identified through either approach were extracted from the annotated genome files (.gbff) based on locus tags and subsequently aligned against the *E. faecalis* OG1 reference locus (NCBI accession no. AE016830.1) using Clinker v0.0.31 (Gilchrist and Chooi 2021). Regions producing alignments (≥30% amino acid identity across individual coding sequences) were classified as ebp-like clusters, whereas regions that did not produce alignments were classified as putative sortase-dependent pilus clusters. Gene clusters aligned using Clinker. All groups are provided in Supplementary Table S3.

### Fixed cell interaction assay

Exponential-phase cultures of OG1RF were normalized to an OD of 0.70 and 1% paraformaldehyde was added. Cultures shook aerobically at 37°C and 200rpm for 16 hours. Samples were aliquoted into microfuge tubes and centrifuged at 94xg. This speed was chosen to prevent damage to pili. Cells were washed 3 times to minimize residual PFA. Fixed OG1RF samples were selectively plated to confirm no viable cells remained. Live *C. difficile* was cocultured with live OG1RF as previously described. Mid-log live *C. difficile* was co-inoculated with fixed OG1RF at 1:10 inoculum for fixed strains and 1:50 inoculum for *C. difficile*. The final wash of OG1RF fixed cultures was added to *C. difficile* as a control for residual PFA.

**Figure S1.**
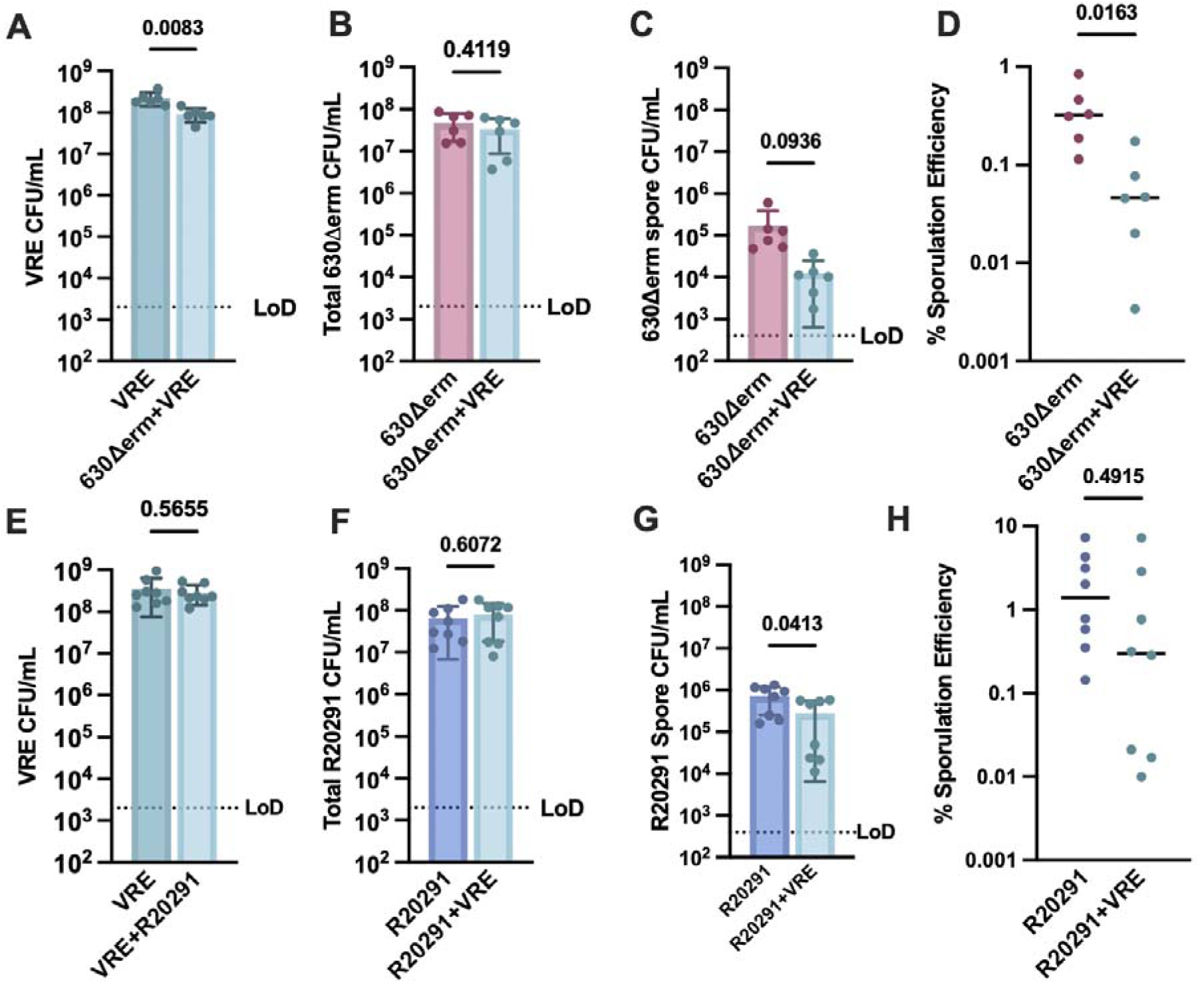
Inibition of sporulation phenotype is somewhat conserved in *C. difficile* strains 630Δ*erm* and R20291. Either R20291 or 630Δ*erm* was cocultured in liquid SMC anaerobically for 48 hours with VRE (A-D) Monoculture vs coculture 630Δerm CFUs with VRE followed by plating on selective media. (A) VRE CFUs (B) Total 630Δerm CFUs (C) Heat-resistant 630Δ*erm* spore CFUs (D) Sporulation efficiency. n = 6 biological replicates carried out in 6 independent expeirments. Statistics: Unparied t-test with Welch’s Correction. (E-H) Monoculture vs coculture of strain R20291 CFUs with VRE followed by plating on selective media. (E) VRE CFUs. (F) Total R20291 CFUs. (G) Heat-resistant R20291 spore CFUs (H) Sporulation efficiency. n = 8 biological replicates carried out in 5 independent experiments. Statistics: Unparied t-test with Welch’s Correction. LoD = Limit of Detection

**Figure S2:**
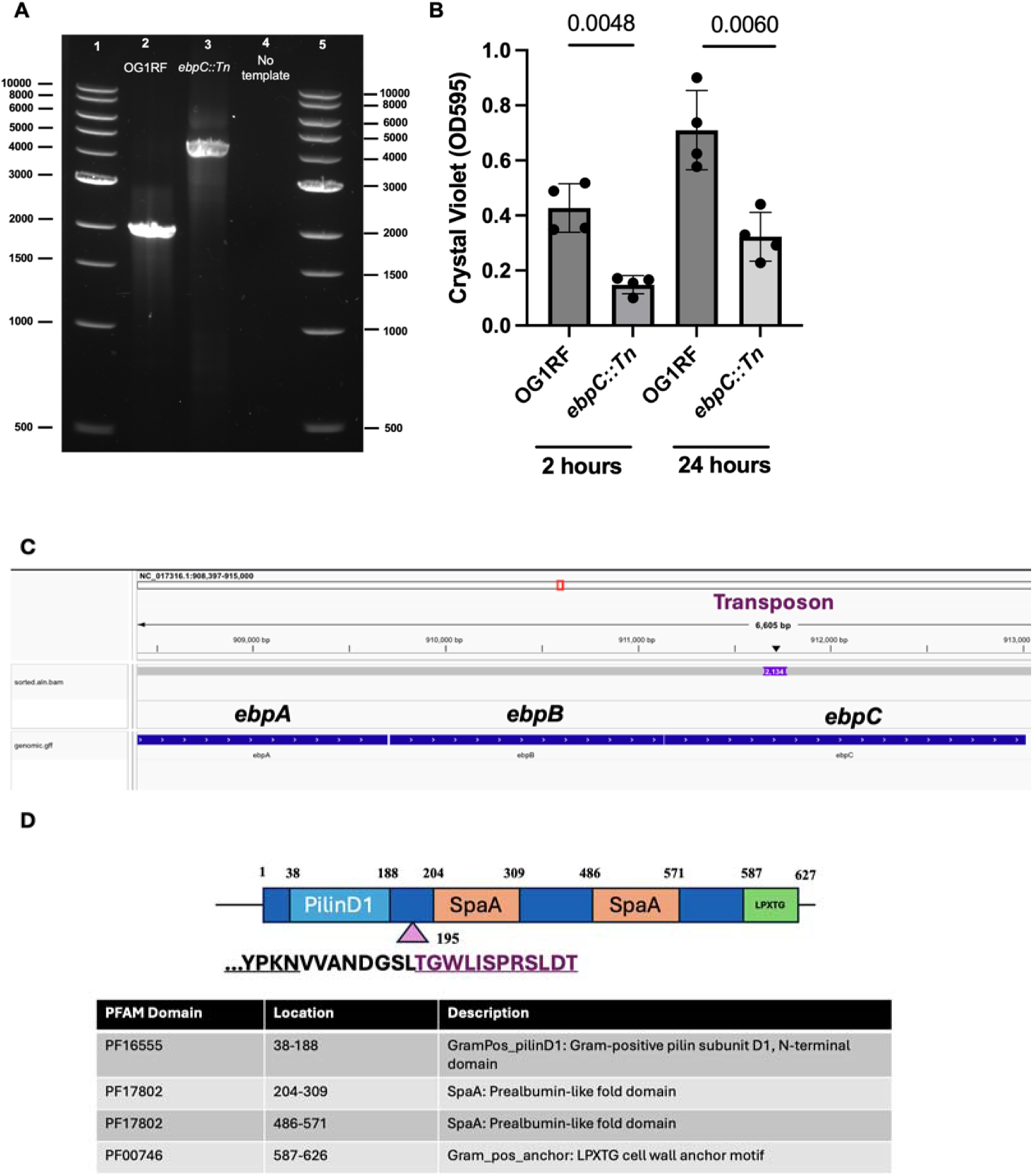
Characterization of *ebpC::Tn*. A) Agarose gel showing amplification of *ebpC* amplified with flanking primers. Lane 1: 1kb Ladder, Lane 2: OG1RF WT template, Lane 3: *ebpC::Tn* template, Lane 4: No template, Lane 5: 1kb Ladder. B) Crystal violet staining of attached cells on a polystyrene plate following 2 hours and 24 hours of incubation. Data are 4 biological replicates. Statistics = unpaired Welch’s t-test. C) Alignment from Integrated Genomics Viewer showing the location of transposon insertion in *ebpC.* D) Diagram of *ebpC* PFAM sequence homology domains. Purple triangle represents transposon insertion site starting at amino acid 195 of the predicted protein sequence. A 12 amino acid portion of the transposon sequence in purple is predicted to be translated before the first in frame stop codon.

**Table S1.**
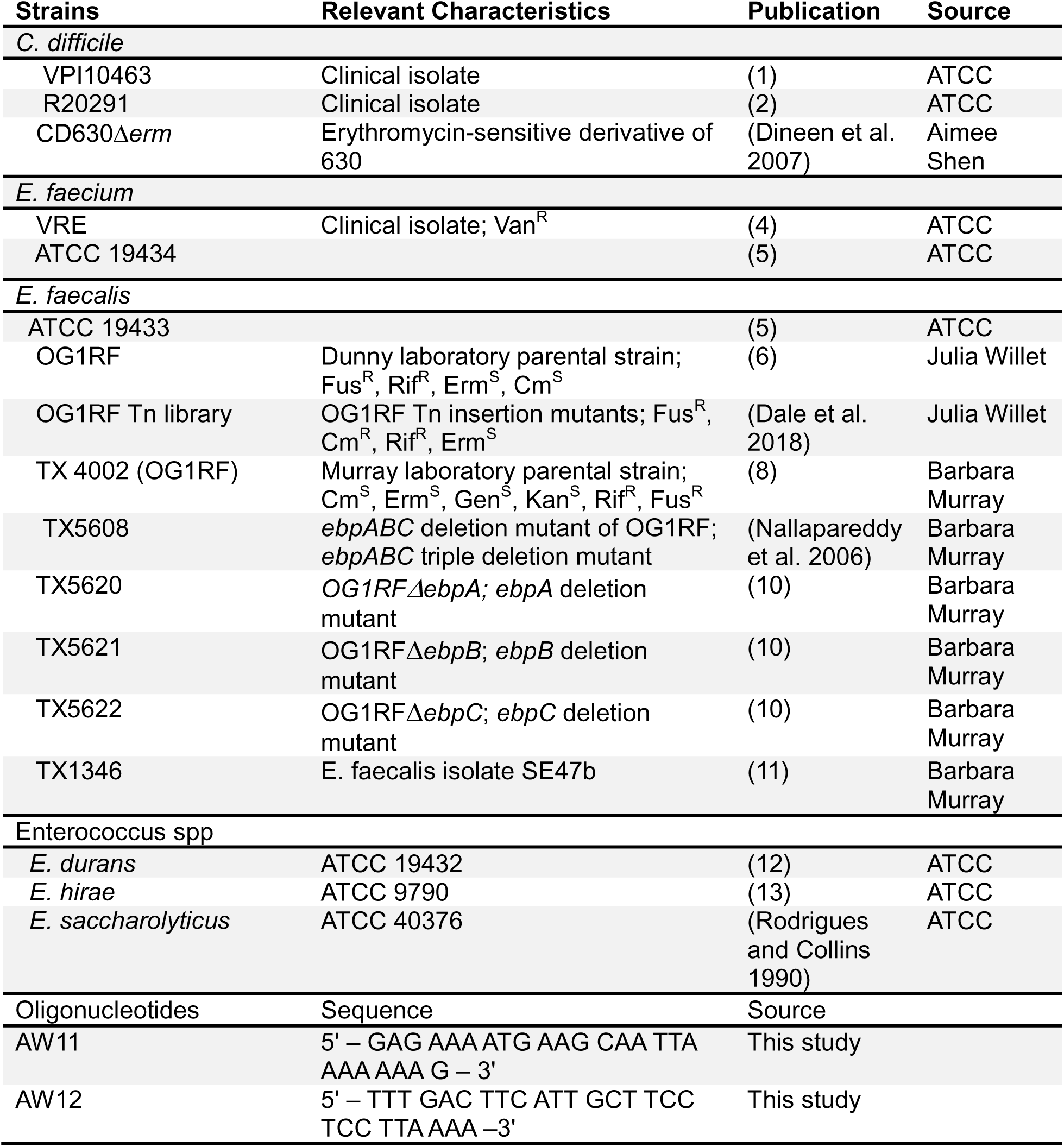

